# Preprocessing Determines Whether Fractal Biomarkers Detect Neurodegeneration: A Systematic Evaluation Across Five fMRI Pipelines

**DOI:** 10.64898/2026.11.06.763877

**Authors:** Tamoghna Chattopadhyay, Harmony Selena Cen, Sophia I. Thomopoulos, Aleksandr Volkov, Yury Maximov, Alan David Motter, Hope Tankesly, Adelaide Hope Ruekgauer, Advait Khopade, Ildar Gabitov, Paul Thompson

**Author notes:** Corresponding author: Tamoghna Chattopadhyay,.

## Abstract

Fractal analysis of resting-state functional MRI (rs-fMRI) has been proposed for detecting altered neural dynamics in Alzheimer’s disease (AD), but the effect of preprocessing on these measurements is poorly constrained. We evaluated 229 ADNI3 participants (156 cognitively normal [CN], 60 mild cognitive impairment [MCI], and 13 AD) processed with five ENIGMA HALFpipe variants. Each scan contained 197 volumes (TR = 3.0 s) and 434 regional time series. The reported CN–AD effect varied 4.5-fold for the detrended fluctuation analysis (DFA) Hurst exponent and 13.1-fold for Sample Entropy; both measures met *p <* 0.05 only after aCompCor. Under aCompCor, mean *H* was 0.812 in CN, 0.827 in MCI, and 0.872 in AD (*d* = −1.40, *p* = 0.0003), whereas the other four DFA effects were approximately *d* = −0.31 to −0.40. A variance decomposition attributed 41.5% of total *H* variance to pipeline, 44.5% to participant, and 0.3% to diagnosis. Whole-brain analysis found higher *H* in AD across 16 of 18 networks, with 36 cortical regions surviving false-discovery-rate correction. Synthetic fractional Gaussian noise experiments confirmed monotonic DFA estimation at the available scan length. The small AD group makes all disease-effect estimates exploratory. These results show that preprocessing choice can determine whether fractal group differences are detectable and should therefore be treated as part of the biomarker definition.

## 1 Introduction

Resting-state BOLD signals exhibit scale-free temporal structure. DFA summarizes this structure with a Hurst exponent, *H*, and has been used to study Alzheimer’s disease, but published results are inconsistent [1]. This inconsistency is methodologically plausible: global signal regression (GSR), censoring, and nuisance regression alter the temporal autocorrelation that fractal estimators are designed to measure. Reviews have therefore called for explicit validation of fractal results against preprocessing choices [2, 3].

We evaluate that sensitivity directly by applying five preprocessing variants to the same 229 participants. Holding the cohort fixed isolates preprocessing from changes in sample composition. The primary question is not clinical classification, but whether common nuisance-control strategies materially change the magnitude and statistical detectability of CN–AD differences in DFA Hurst exponent and Sample Entropy.

## 2 Materials and Methods

### 2.1 Data and Preprocessing

The ADNI3 sample comprised 156 CN, 60 MCI, and 13 AD participants. Each rs-fMRI series contained 197 BOLD volumes at TR = 3.0 s (approximately 10 minutes). Signals were parcellated into 434 regions using the Schaefer-400 cortical atlas plus 34 subcortical and cerebellar regions.

The same participants were processed with five ENIGMA HALFpipe variants: v1, aCompCor nuisance components derived by white-matter/cerebrospinal-fluid PCA; v2, six motion parameters plus scrubbing; v3, motion, scrubbing, and GSR; v4, motion parameters alone; and v5, motion parameters plus GSR. GSR can temporally downweight shared fluctuations, while scrubbing interrupts the contiguous series assumed by DFA [4, 5].

### 2.2 Fractal Measures and Statistics

DFA integrated each regional time series, removed local linear trends, and estimated the slope of log *F* (*n*) against log *n* over 20 log-spaced windows from 4 to *N/*4 samples [6]. Sample Entropy provided a complementary measure of temporal irregularity. Multifractal DFA at *q* = 2 and Higuchi fractal dimension (HFD) were included in the synthetic validation. Group comparisons used Mann–Whitney *U* tests and Cohen’s *d*; regional comparisons used Benjamini–Hochberg false-discovery-rate control at *q <* 0.05. Age sensitivity was evaluated by linear residualization and an age-matched subcohort (±3 years).

Estimator behavior at the available length (*N* = 197) was tested on fractional Gaussian noise with known *H* ∈ { 0.3, 0.5, 0.7, 0.9}, 50 trials per value. DFA showed a positive bias of approximately 0.05–0.09 but remained monotonic; a true Δ*H* = 0.2 produced *d* = 2.62 (Fig. 3).

**Figure 1.**
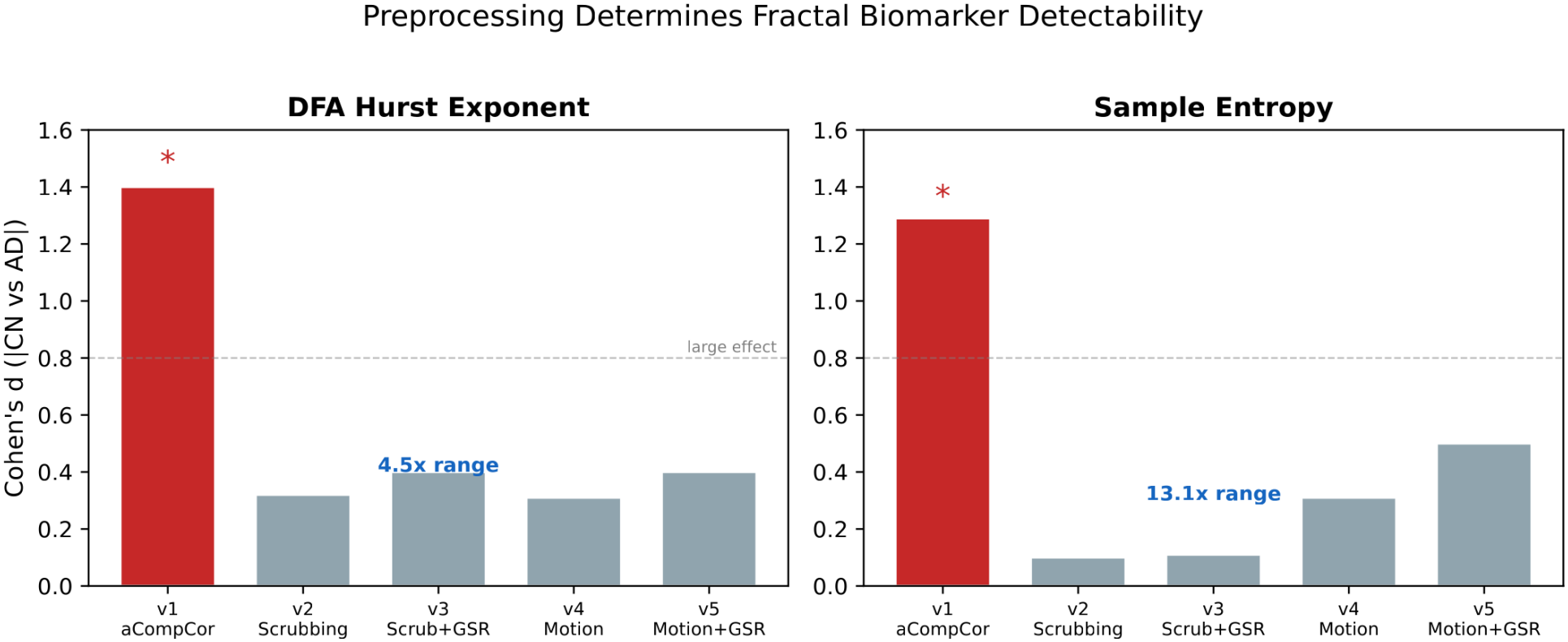
CN–AD effect-size magnitude for DFA Hurst exponent (left) and Sample Entropy (right) across five preprocessing pipelines. Red bars mark *p <* 0.05. The same participants were used in every pipeline; only aCompCor yielded significant effects for both measures.

**Figure 2.**
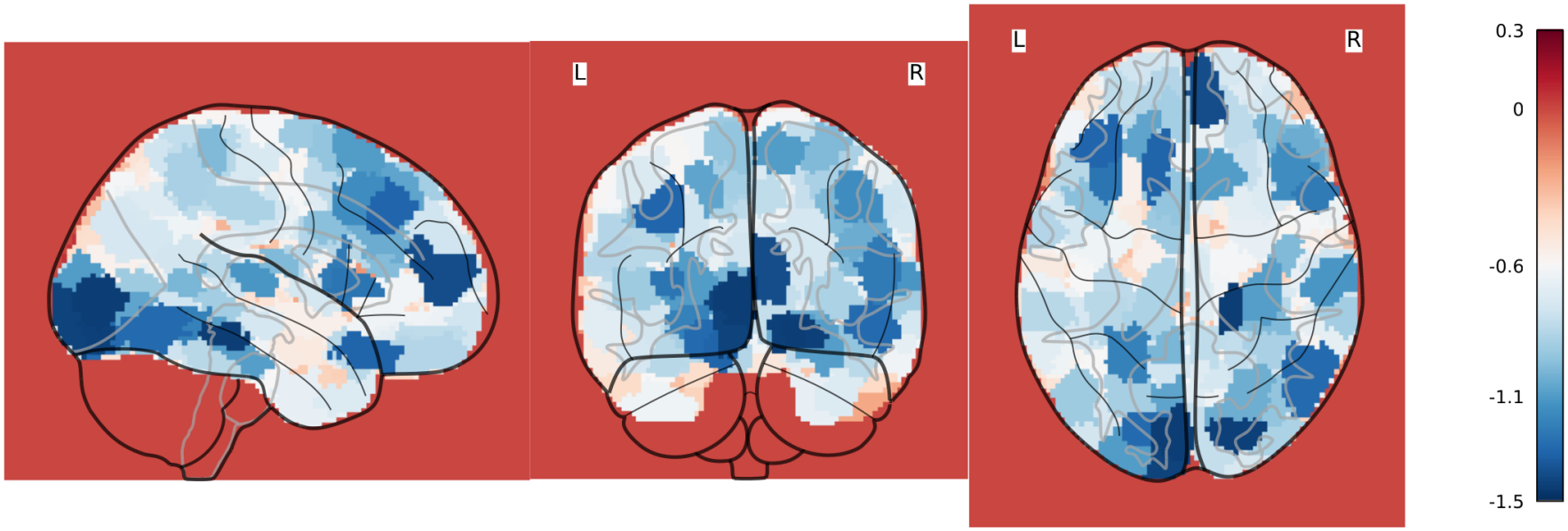
Glass-brain projection of CN–AD DFA Hurst effect sizes across 400 Schaefer cortical regions. Blue indicates higher *H* in AD; effects span prefrontal, parietal, temporal, and occipital cortex.

**Figure 3.**
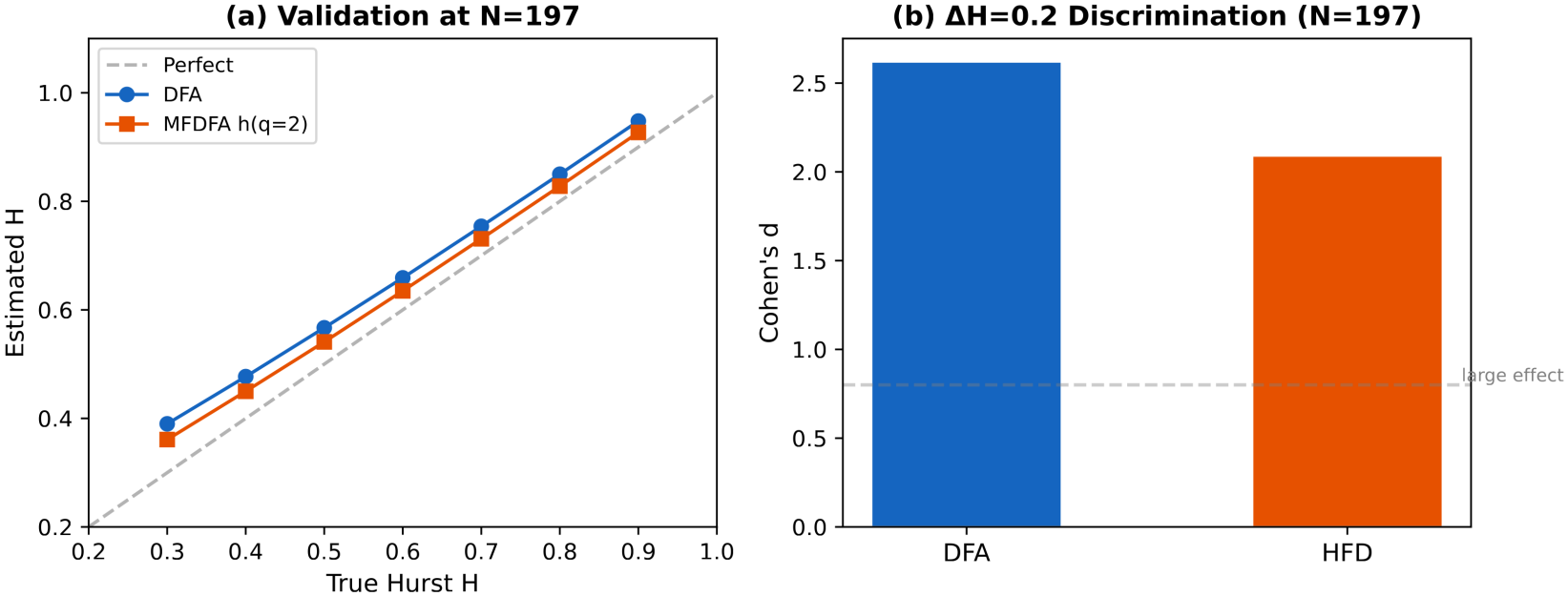
Synthetic validation at *N* = 197. (a) DFA and MFDFA estimates remain monotonic despite positive bias. (b) A true Δ*H* = 0.2 produces large standardized separation for DFA and HFD.

## 3 Results

### 3.1 Pipeline Sensitivity

The absolute Hurst level fell from approximately 0.81 after aCompCor to approximately 0.67 in the GSR variants. More importantly, the reported CN–AD DFA effect varied from *d* = −1.40 after aCompCor to *d* = −0.31 to −0.40 after the alternatives, a 4.5-fold range (Fig. 1). Only aCompCor met the reported *p <* 0.05 threshold for DFA (*p* = 0.0003). Sample Entropy was even more pipeline-sensitive: its effect range was 13.1-fold, and again only aCompCor was significant (*d* = 1.29, *p* = 0.0005). Cross-pipeline participant correlations were lower between aCompCor and the other variants (*r* = 0.55) than among v2–v5 (*r* = 0.75–0.98).

A decomposition of all 1,145 subject–pipeline observations attributed 41.5% of total *H* variance to preprocessing pipeline and 44.5% to participant, compared with 0.3% to diagnosis. Thus, pipeline-associated variance was approximately 120 times the diagnosis-associated component in this analysis.

### 3.2 Spatial Distribution and Age Sensitivity

Across 434 regions, 16 of 18 functional networks showed higher *H* in AD, and 36 regions survived FDR correction. The strongest reported network effects occurred in visual cortex (*d* = − 1.45), subcortical structures (*d* = − 1.10), and default-mode cortex (*d* = − 1.04). The cortical projection (Fig. 2) shows that the effect was spatially distributed rather than confined to one region.

AD participants were 8.0 years older than CN participants. The pooled age–*H* association was weak (*r* = 0.095, *p* = 0.15). The CN–AD difference persisted after linear age residualization (*d* = −0.78, *p* = 0.019) and in an age-matched subcohort (*d* = −0.86, *p* = 0.008), although neither analysis removes all possible age, site, scanner, or motion confounding.

## 4 Discussion

Applying five pipelines to identical participants shows that preprocessing is not a neutral prelude to fractal analysis; it helps define the quantity being measured. aCompCor retained a large CN–AD contrast for both DFA and Sample Entropy, whereas GSR and scrubbing strongly attenuated statistical detectability. The variance decomposition and reduced cross-pipeline correlation with aCompCor reinforce that this is more than a shift in the group mean.

The results support routine multi-pipeline sensitivity analysis, complete reporting of censoring and nuisance regression, and caution when comparing absolute *H* across studies. They do not by themselves prove that aCompCor recovers a purely neural signal or that GSR and scrubbing remove biology. GSR targets shared low-frequency variance, and scrubbing changes temporal continuity; both operations directly interact with DFA’s estimand. Physiological recordings and an independent disease cohort are needed to distinguish biological recovery from residual-noise effects.

The present analysis is limited by the relatively small Alzheimer’s disease cohort, motivating ongoing expansion of the dataset to improve statistical power and the stability of estimated disease effects. The spatial analysis also entails many regional tests despite FDR control. These results should therefore be interpreted as an evaluation of measurement sensitivity, not evidence of diagnostic performance. A confirmatory analysis should enlarge the AD group, lock endpoints before inspection, and model diagnosis-by-pipeline effects with participant as the repeated-measures unit.

### 4.1 Future Work

Ongoing work will extend this evaluation using additional ADNI dementia participants that have recently completed preprocessing and quality control. Incorporating these new subjects will substantially increase the Alzheimer’s disease cohort and enable a more robust assessment of preprocessing-dependent fractal biomarkers. Future analyses will re-evaluate effect sizes, variance decomposition, and regional findings in the expanded dataset, while further examining the influence of acquisition site, motion, and other potential confounds on biomarker stability.

## 5 Conclusion

Across five preprocessing variants applied to the same participants, fractal CN–AD effects ranged 4.5-fold for DFA and 13.1-fold for Sample Entropy and were significant only after aCompCor. Preprocessing sensitivity should therefore be evaluated and reported as part of any fractal fMRI biomarker claim.

## Acknowledgments

Data used in preparation of this article were obtained from the Alzheimer’s Disease Neuroimaging Initiative (ADNI) database (adni.loni.usc.edu). As such, the investigators within the ADNI contributed to the design and implementation of ADNI and/or provided data but did not participate in the analysis or writing of this report. A complete listing of ADNI investigators can be found in the [ADNI investigator acknowledgment list](https://adni.loni.usc.edu/wp-content/uploads/ADNI%20Acknowledgement%20List_Feb2025_clean.pdf).

The ADNI imaging and clinical data came from the [Alzheimer’s Disease Neuroimaging Initiative database](https://adni.loni.usc.edu/). ADNI began in 2003 as a public-private partnership led by Michael W. Weiner, MD. It brings together longitudinal imaging, biomarkers, and clinical and neuropsychological assessments to study progression from mild cognitive impairment to Alzheimer’s disease.

The National Institute on Aging funds ADNI data collection and sharing through National Institutes of Health grant U19 AG024904 to the Northern California Institute for Research and Education. ADNI has also received support from the National Institute of Biomedical Imaging and Bioengineering, the Canadian Institutes of Health Research, and private contributors through the Foundation for the National Institutes of Health. These contributors include AbbVie, Alzheimer’s Association, Alzheimer’s Drug Discovery Foundation, Araclon Biotech, BioClinica, Inc., Biogen, Bristol-Myers Squibb Company, CereSpir, Inc., Cogstate, Eisai Inc., Elan Pharmaceuticals, Inc., Eli Lilly and Company, EuroImmun, F. Hoffmann-La Roche Ltd and its affiliated company Genentech, Inc., Fujirebio, GE Healthcare, IXICO Ltd., Janssen Alzheimer Immunotherapy Research & Development, LLC., Johnson & Johnson Pharmaceutical Research & Development LLC., Lumosity, Lundbeck, Merck & Co., Inc., Meso Scale Diagnostics, LLC., NeuroRx Research, Neurotrack Technologies, Novartis Pharmaceuticals Corporation, Pfizer Inc., Piramal Imaging, Servier, Takeda Pharmaceutical Company, and Transition Therapeutics.

